# Structural changes after early life adversity in rodents: a systematic review with meta-analysis

**DOI:** 10.1101/2022.01.17.476580

**Authors:** Marian Joëls, Eline Kraaijenvanger, R. Angela Sarabdjitsingh, Valeria Bonapersona

**Author notes:** Corresponding Author: (VB).

## Abstract

Early life adversity (ELA) is a well-documented risk factor for psychiatric illnesses in humans. This risk may, in part, be conferred by structural changes induced by ELA, lasting into adulthood. We here review the evidence for such lasting structural changes in rodent models for ELA involving altered maternal care during the first two postnatal weeks. In total, we extracted data from 64 studies reporting on 260 comparisons in adult rats or mice which experienced ELA or control treatment. Most of the observations concerned structural changes in the hippocampus of adult male rats earlier exposed to maternal separation. A 3-level meta-analysis revealed that ELA reduced hippocampal volume and the number of dendritic branches as well as dendritic length of principal hippocampal cells. No differences were observed across the hippocampal subfields. In terms of adult neurogenesis in the dentate subgranular zone, both staining for BrdU and the early neuronal marker DCX were significantly reduced, while the general proliferation marker Ki67 remained unchanged. The neuronal growth factor BDNF did not show significant changes, although the unexplained heterogeneity was moderate. Generally, the effect of ELA compared to control on structural markers was not affected by additional stressors experienced in life. Overall, the data available support the notion that ELA, at least in the hippocampus of male rats, lastingly reduces volume, hampers dendritic growth and suppresses adult neurogenesis.

## Introduction

Adversity experienced early in life, when the brain is still developing, is a well-documented trigger for lasting changes in brain connectivity and behavior in humans (Chen & Baram, 2016; Herzberg & Gunnar, 2020; Sandi & Haller, 2015; Teicher et al., 2016, 2021), presumably increasing the risk of individuals to develop psychopathology later in life (Brown et al., 2019; Hughes et al., 2017). To study the long-lasting consequences of early life adversity (ELA) on brain structure and function, researchers have often reverted to animal models (Benmhammed et al., 2019; Branchi & Cirulli, 2014). These offer multiple advantages over human investigations, e.g. that i) early life environment is known and can be specifically altered; ii) genetic variation, especially in in-bred mice, is relatively low; iii) housing conditions can be kept constant; and iv) the lifespan is quite short, so that consequences of ELA for the adult brain can be studied over the courses of months rather than decades (Knop et al., 2017).

Recently, we and others have shown in meta-analyses that in animal models too, ELA results in very consistent changes in behavioral function (Bonapersona et al., 2019; Rocha et al., 2021; D. Wang et al., 2020). Shifts in behavior likely result from alterations in the underlying neuronal substrate. This can relate to many ELA-induced changes, including the connectivity between areas, structural modifications within specific areas and cell types, but also functional changes related to neurotransmitter actions and/or the cellular responses downstream of neurotransmitters and their receptors, such as second messenger systems or gene transcription. Also, the response of the individual animal to stressful circumstances may be altered in ELA-exposed compared to control animals (Anacker et al., 2014), which may affect behavioral outcome particularly in challenging tasks, e.g. contextual fear conditioning.

To date, the effects of ELA on neuronal structure and structural plasticity in adult rodents have not been examined meta-analytically. We here focused on studies in rats and mice describing the effects of postnatal adversity (i.e., starting during the first 2 weeks of life) on structural outcome; all models involved altered maternal care, an important environmental determinant of adversity during this developmental window. We focused on three sets of outcomes: First, volume of adult brain areas and morphology of neurons after ELA, including reports on dendritic tree morphology and spine density. Secondly, adult neurogenesis in the dentate gyrus (Kempermann et al., 2015) of animals with an ELA history, summarizing data on cell proliferation, differentiation and survival. And thirdly, studies reporting on brain derived nerve growth factor (BDNF), which might give insight in potential mechanisms by which structure could be changed. For the main analysis, a 3-level mixed effect model was applied (Bonapersona et al., 2018; Cheung, 2014). In case of significant unexplained heterogeneity, we next performed an exploratory random forest analysis (Lissa, 2020) to identify the most promising potential moderators. Since the effects of stress have often been shown to differ between males and females (Bale & Epperson, 2015; Bangasser & Cuarenta, 2021), we planned in advance to conduct our analyses for males and females separately.

## Methods

This review adheres to SYRCLE’s guidelines for protocol (De Vries 2015), search strategy (Leenaars et al., 2012), and risk of bias assessment (Hooijmans et al., 2014). Reporting is in accordance with the PRISMA reporting checklist (Moher et al., 2009, **Supplementary Information S2).** The analytic strategy is based on earlier work of our own lab (Bonapersona et al., 2018, 2019). Materials, data and scripts used for this project are available via the open science framework (*https://osf.io/9gru2/*).

### Search strategy and data gathering

To investigate the effects of ELA on structural plasticity, we conducted a systematic literature search on April 3^rd^ 2019 on the electronic databases PubMed and WebOfScience. The search string included the terms ‘mice and rats’ and ‘postnatal ELA’ **(Supplementary note** 1), which was previously already used by our own lab (Schuler et al., 2021). For this particular study, ELA was defined as all postnatal models that are based on alterations in maternal care (Levine, 2002), either experimentally induced (through maternal deprivation of pups (Marco et al., 2015); separation or isolation of dam and pups; or exposing dams and their litter to limited nesting and bedding material (Rice et al., 2008)), or naturally (i.e. variations in the amount of licking and grooming of pups by the dam (Champagne et al., 2003), for a review (Benmhammed et al., 2019)). Study selection was performed in Rayyan (Ouzzani et al., 2016) by at least two researchers (see also acknowledgements, HS, DvN, LvM). The order in which the publications were assessed differed across researchers, and it occurred in two stages. In the first stage, studies were excluded based on titles and abstracts if they: 1) were not a primary publication, 2) did not use mice or rats, 3) were not related to early life adversity. During the second stage, full text was screened and studies were included if: 1) structural plasticity outcomes were measured; 2) the outcomes were measured in adulthood (older than 8 weeks but younger than 1 year of age); 3) the animals and previous generations did not experience other pharmacological / dietary / genetic interventions; 4) the animals were not germ free, were not specifically bred for certain traits and were not reported in split groups (e.g. high/low performance); 6) the sex of animals was known (either based on the report or after contacting authors); 7) in the intervention models, the control group was separated from the mothers for less than 5 min (i.e. the “handling” model was excluded). Disagreements in study selection were resolved by involving an independent scientist (MJ). An overview of the study procedure is shown in **Figure 1**.

**Figure 1.**
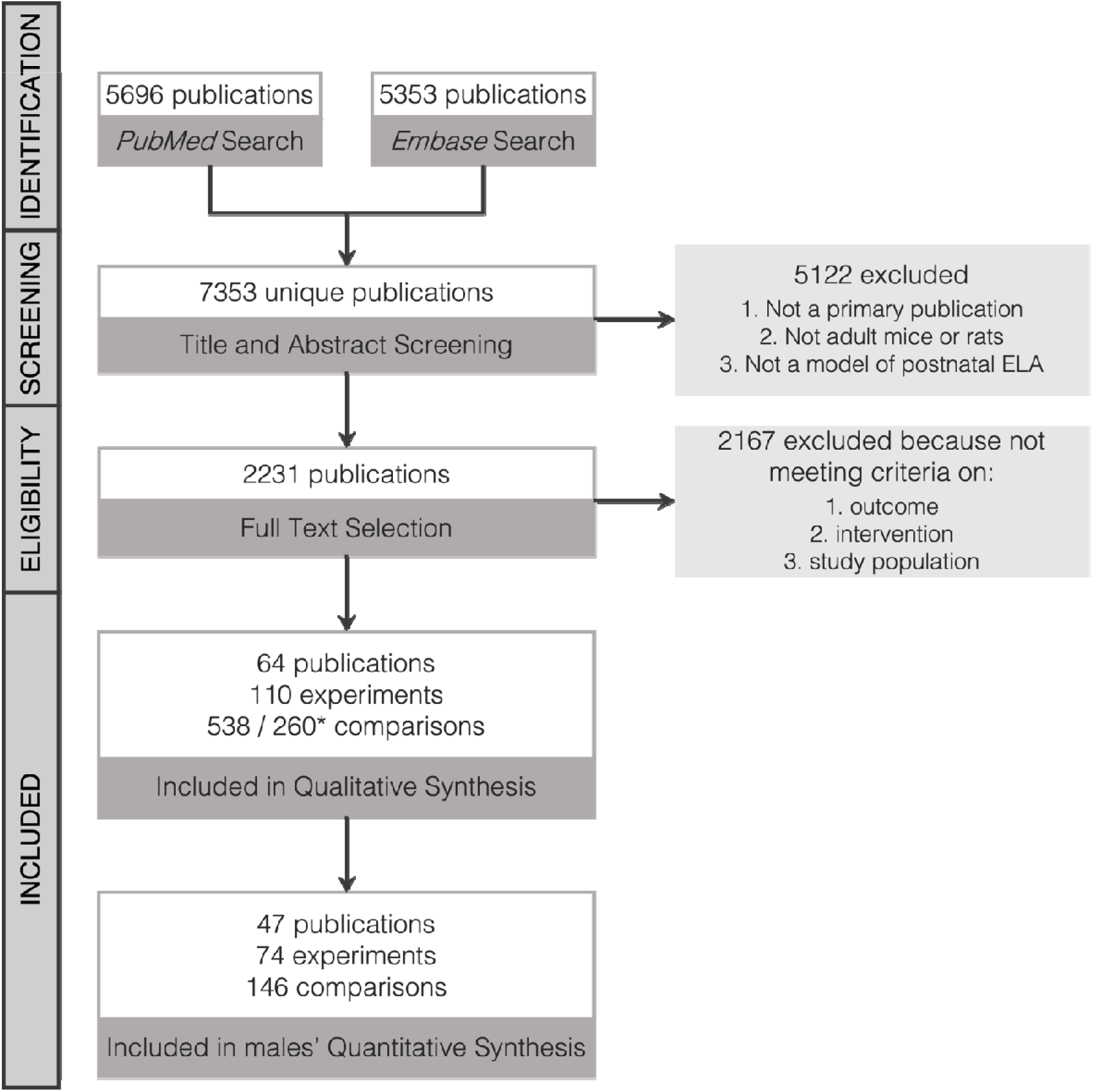
PRIMA flowchart for study selection and inclusion. * = 538 is the number of comparisons before preprocessing; 260 refers to the number included after processing. For information about preprocessing, see *Methods*.

Data from eligible studies was organized in a standardized database, which is available via the open science framework (*https://osf.io/9gru2/*). Two reviewers (VB and EK) shared the data collection task. Papers considered unclear were evaluated by both reviewers independently, and subsequently discussed with a third reviewer (MJ). It includes details about 1) the publications (author, year, reference), 2) the experimental design (e.g. species, sex, model, other life events, age and state of the animals at the time of testing), 3) information about the outcome extracted (e.g. brain area, technique used), and 4) summary statistics of the data measured (e.g. sample size, mean or median, and deviation or interquantile range (IQR)). According to this structure, we organized the information of each individual comparison between a control and an experimental group. Of note, the groups always differed only in the experience of ELA. All other variables (e.g. additional ‘life events’) were comparable between the control and ELA group.

If only SEM was reported, SD was calculated as SEM * n, where n is the number of animals per group. If median and IQR were reported rather than mean and SD, we evaluated whether the median could be an approximation of the mean, i.e. the median was in the approximate center of the IQR range. If this condition was not met (n_cmp excluded_ = 2), the comparison was excluded as it was not possible to obtain an effect size measure comparable to that of the other publications. If the condition was met, the median was transformed to mean according to (Hozo et al., 2005) formula:

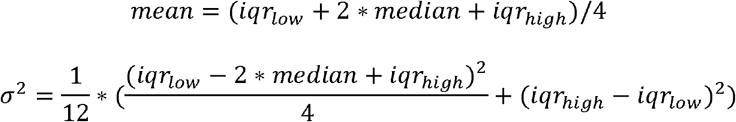

If the number of animals were reported as a range (e.g. 6–8 animals per group), we used the lower boundary of this number (e.g. 6 animals per group) as a conservative estimate. Data that was reported exclusively in graphs was digitalized with WebPlotDigitizer (Rohatgi, Ankit, 2021). For all remaining missing information, we contacted the corresponding author of each manuscript after 2008 (response rate 80%). If no answer was received after 2 months and a reminder, we considered the data as not retrievable.

#### Data synthesis and statistical analysis

Effect sizes for each individual comparison (i.e. the standardized mean difference between control and ELA on each specific outcome) were calculated with *escalc* (R package *metaphor*, version 3.0-2 (Viechtbauer, 2010)) using the Hedge’s g (g) method, which includes a correction factor for small sample sizes (Vesterinen et al., 2014).

For the main analysis, we used a 3-level mixed effect model which accounts for the anticipated heterogeneity of the studies as well as the dependency of effects within experiments (Cheung, 2014). In our experimental design, the 3 levels correspond to variance of effect size between 1) animals, 2) outcomes and 3) publications. Structural plasticity was broadly classified in three sets of outcome: a) (neuronal) morphology, b) neurogenesis, and c) growth factors, specifically BDNF. Given their different biological meaning, these were analyzed in separate models. Prior to the start of the study, we defined potential moderators of the effects of ELA on structural plasticity, namely: 1) specific outcome parameters, 2) brain area(s), 3) experience of other traumatic events, 4) product measured (mRNA or protein, only for the outcome BDNF), 5) state of the animal at death (only for BDNF and neurogenesis, i.e. rest or not), 6) delay between the start of the experimental manipulation and measuring the outcome (only for the neurogenesis-related parameter BrdU where delay between injection and measurement gives rise to a specific interpretation of the data). The final moderators were selected based on the frequency of the available literature, to maximize interpretability and robustness of the results. Supplementary Table 1 summarizes the final models and the considerations taken.

Since most of the analytical decisions were based on frequencies, some categorizations were modified after data collection (but before analysis). These changes were based uniquely on the frequencies, with the intent to maximize the balance between subcategories, while maintaining interpretability. Generally, we used as a rule of thumb that a category could be analyzed only if it was investigated by at least 4 independent publications. Categorizations were therefore adapted to meet this requirement. Specifically, the state of the animals at death was initially coded as rest, aroused or stressed. However, due to the limited number of comparisons in the aroused/stressed categories, these were merged into a “not rest” category. Furthermore, if a study reported multiple sub-brain areas within one of our categorizations, these were combined for the quantitative synthesis to limit heterogeneity and over representation of a certain outcome in the analysis. Similarly, if a study reported multiple outcomes (e.g. multiple BDNF exons), these were combined into one measure. For volumes, this was achieved by adding together summary statistics. Given 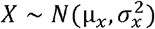 and 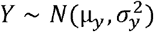, *Z* = *X* + *Y*. Then, 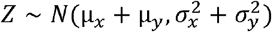

For all other outcomes, effect sizes were merged with the *metafor*’s function *aggregate* (Viechtbauer, 2010). For all analyses, p-values were adjusted for multiple comparisons using the *Holm* correction (Holm, 1979).

Lastly, we performed an additional exploratory subgroup analysis to compare specifically basal vs apical dendrites in the CA1 and CA3 hippocampal areas. In the main analysis, basal and apical dendrites were merged together in a unique effect size. For this analysis, we build two 3-levels mixed effect models (one for basal and one for apical) and compared them with a Wald test. Of note, we used all publications reporting on either basal and apical, because only 2 publications reported both within the same animals.

Heterogeneity was tested with Cochran Q-test (Higgins & Thompson, 2002) and I^2^ statistics (Cheung, 2014). A significant test of heterogeneity or a large I^2^ (see rules of thumb in (Cheung, 2014)) signifies that the data still has variance that cannot be explained by chance alone, despite the used moderators. For those models with significant unexplained heterogeneity (i.e., neurogenesis and BDNF), we next performed an exploratory analysis to explore the source of the unexplained heterogeneity. This was conducted using *MetaForest* (van Lissa, 2020), a novel exploratory approach to identify the most promising potential moderators to explain heterogeneity. This method is an application of random forests to meta-analysis data by means of bootstrap sampling. *MetaForest* ranks moderators based on their (non)linear influence on the effect size. Here, this analysis was conducted for neurogenesis and BDNF separately. We selected 15 potential moderators for *Metaforest* analysis. After identifying a convergence range, we conducted a recursive preselection based on 100 replications, and selected only those moderators that were selected in at least 50% of the replications. With these variables, we built our *MetaForest* model and conducted a 10x cross validation to determine the optimal tuning parameters that minimized root-mean-square deviation (for neurogenesis: random weights, 2 candidate moderators at each split, minimum node size =4; for BDNF: fixed weights, 4 candidate moderators at each split, minimum node size =5). To estimate how much variance was explained by our model, we calculated the cross-validated R^2^(R_cv_^2^), which is robust to overfitting and provides evidence for the results’ generalizability.

#### Sex differences

Prior to the beginning of the study, we planned to conduct our analyses for males and females separately, since the effects of stress on brain and behavior have often been shown to differ across sexes (Bale & Epperson, 2015; Bangasser & Cuarenta, 2021), at least regarding effect size (Bonapersona et al., 2019). However, due to the limited number of publications in females, quantitative analysis was feasible only in the males’ dataset. As an alternative, we focused on investigating sex differences in a subset of publications reporting data on both sexes. Although with this sex-matched dataset it still was not possible to explore sex differences related to specific outcomes, we can investigate whether there are fundamental sex differences in the effect sizes, for example due to male-developed ELA models (Bonapersona et al., 2019).

We calculated the effect sizes (Hedge’s *g*) for males and females separately on a subset of studies, i.e. the sex-matched dataset. Two identical models were built for the data subsets, one for each sex, without moderators due to the limited amount of evidence available. In these models, we used the absolute value of the effect size since the different outcomes may have opposing effects thereby cancelling each other out in the meta-analytic model. We then performed a Wald test to compare the female versus male models.

#### Bias assessment

To assess risk of bias, we followed SYRCLE’s risk of bias guidelines (Hooijmans et al., 2014). Two reviewers (EK and VB) assessed risk of bias independently on the whole dataset, and resolved disagreements with discussion. To the best of our knowledge no quantitative method is available for the inspection of publication bias for a multi-level setting. Publication bias was therefore assessed on the univariate models for each of the functional outcomes (morphology, neurogenesis and BDNF) by qualitative inspection of funnel plot asymmetry, adapted using a measure of pooled standard deviation in the formula for precision (1/variance) as suggested by Vesterinen and colleagues (Vesterinen et al., 2014). Contrary to our initial study protocol, we did not conduct a Egger’s regression, because the number of publications was low and Egger’s regression would have most likely been underpowered (Sterne et al., 2011). Rather, we interpret the probable influence of publication bias based on the areas of significance, following (Sterne et al., 2011).

#### Software

The analyses were conducted in R (version 3.5.1) (R Core Team, 2015), using the following packages: 1) *metafor* version 3.0-2 (Viechtbauer, 2010) for conducting the analysis, 2) *metaforest* version 0.1.3 (Lissa, 2020) for data exploration, 3) *dplyr* version 1.0.7 (Wickham et al., 2021) for general data handling. The R script and data are available (*https://osf.io/9gru2/*).

## Results

### Study selection and qualitative analysis

An overview of the study design is summarized in the flow chart **(Figure 1**). Our pre-specified inclusion criteria (see Methods) were met by 64 publications, published between 2002 and 2018. The included publications contributed 110 unique experiments, with a total of 260 comparisons from which we extracted statistical measurements (e.g. mean, standard deviation (SD) and sample size (N)). 9 comparisons from 3 publications were excluded from the analysis, as it was not possible to extract nor infer any statistical measurement.

The included publications used mainly rats (n_publ_ = 83%) and the maternal separation model to mimic ELA (n_publ_ = 66%), followed by the limited nesting and bedding model (n_publ_ = 17%), maternal deprivation (n_publ_ = 11%), observation of natural variations in licking and grooming (n_publ_ = 5%) and isolation (n_publ_ = 1-2%).

To study structural plasticity after ELA, we focused on morphology (i.e., changes in the size of brain areas, morphology of dendrites or spine density), neurogenesis (i.e., staining for BrdU, DCX and Ki67) and the growth factor BDNF. **Table 1** summarizes the frequency of each outcome.

**Table 1.** Outcome frequencies in both sexes. The functional categorizations correspond to the classifications for the analyses. # = number

| Functional interpretation | outcome | #studies | #experiments | #comparisons |
| --- | --- | --- | --- | --- |
| morphology | volume | 10 | 17 | 35 |
| morphology | dendritic changes | 14 | 17 | 49 |
| morphology | spines | 6 | 8 | 18 |
| neurogenesis | BrdU + cells | 16 | 28 | 30 |
| neurogenesis | DCX staining | 13 | 17 | 17 |
| neurogenesis | Ki67 staining | 11 | 14 | 14 |
| growth factor | BDNF | 28 | 44 | 97 |

In total, more than 10 brain areas were investigated, with most studies describing the hippocampus (67% of all comparisons in males and females, **Figure 2)**. Within the hippocampus, 81 comparisons were from the dentate gyrus, 22 from the CA1, 22 from the CA3, 3 from the CA4, while 47 measured the whole hippocampus (unspecified); CA4 was excluded from further analysis because of a too low number.

**Figure 2.**
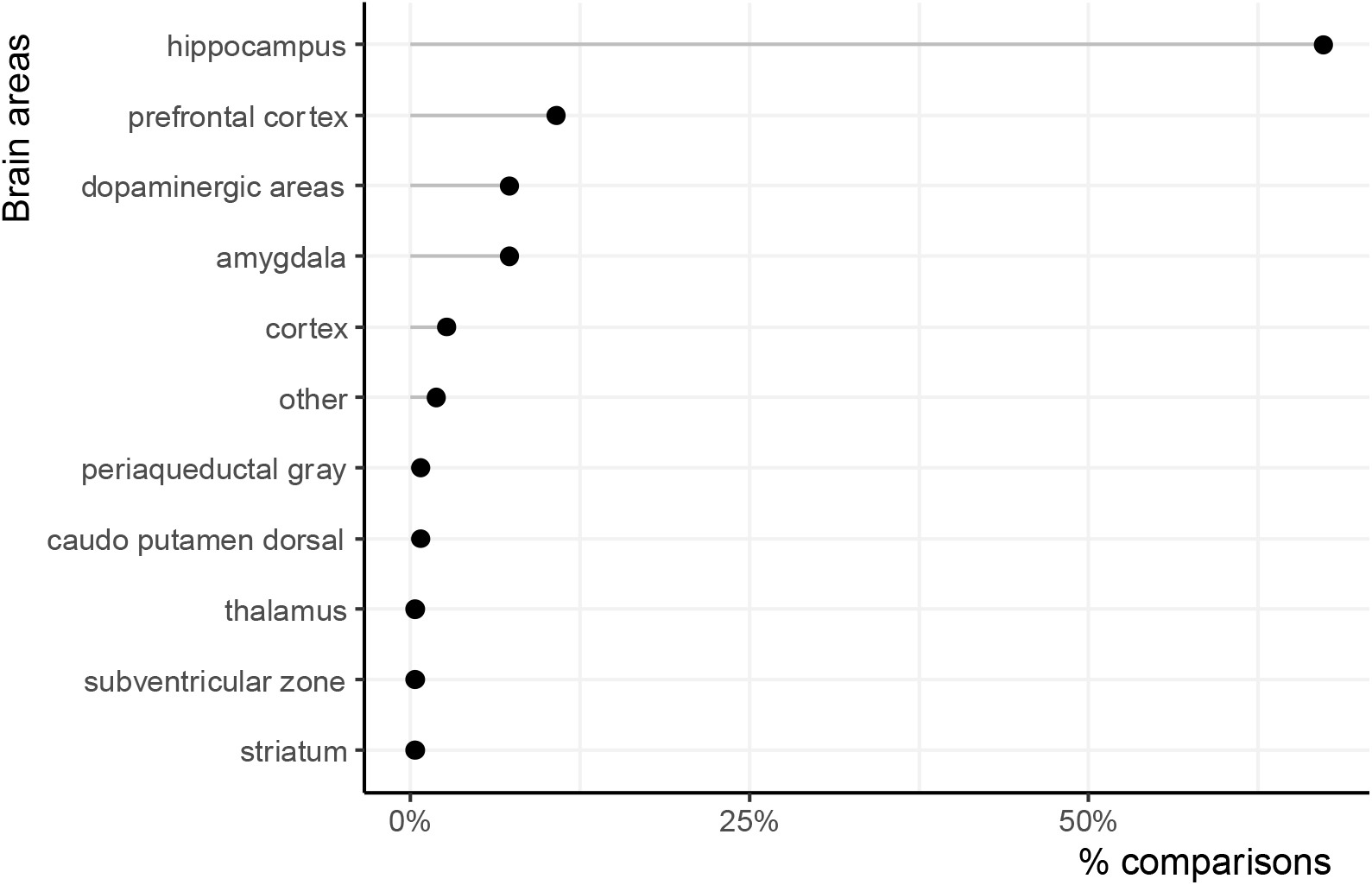
Distribution of the brain areas investigated in relationship to ELA and structural plasticity. The categorizations follow the Allen Brain Atlas embryological classification.

A total of 3336 animals were used, of which the majority (79%) was male. Thirteen publications performed experiments in both male and female rodents. Similar to previous studies, we aimed to analyze males and females separately, i.e. as two different biological systems, since sex-dependent characteristics have been frequently observed in stress research (Bale & Epperson, 2015; Bangasser & Cuarenta, 2021). However, data on females was too scarce to be analyzed quantitatively at a meta-level. We therefore focused our quantitative analyses on males, and subsequently performed an exploratory analysis on a subset of the data to explore potential sex differences.

Based on the frequencies reported above, we included only the outcomes of the hippocampus in subsequent quantitative synthesis, in male mice. Similarly, the number of dendrites (n_publ_ = 2) and spine density (n_publ_ = 3) were not included in the meta-analyses due to the limited number of publications. The descriptive results of these two parameters are summarized in **Table 2**.

**Table 2.**
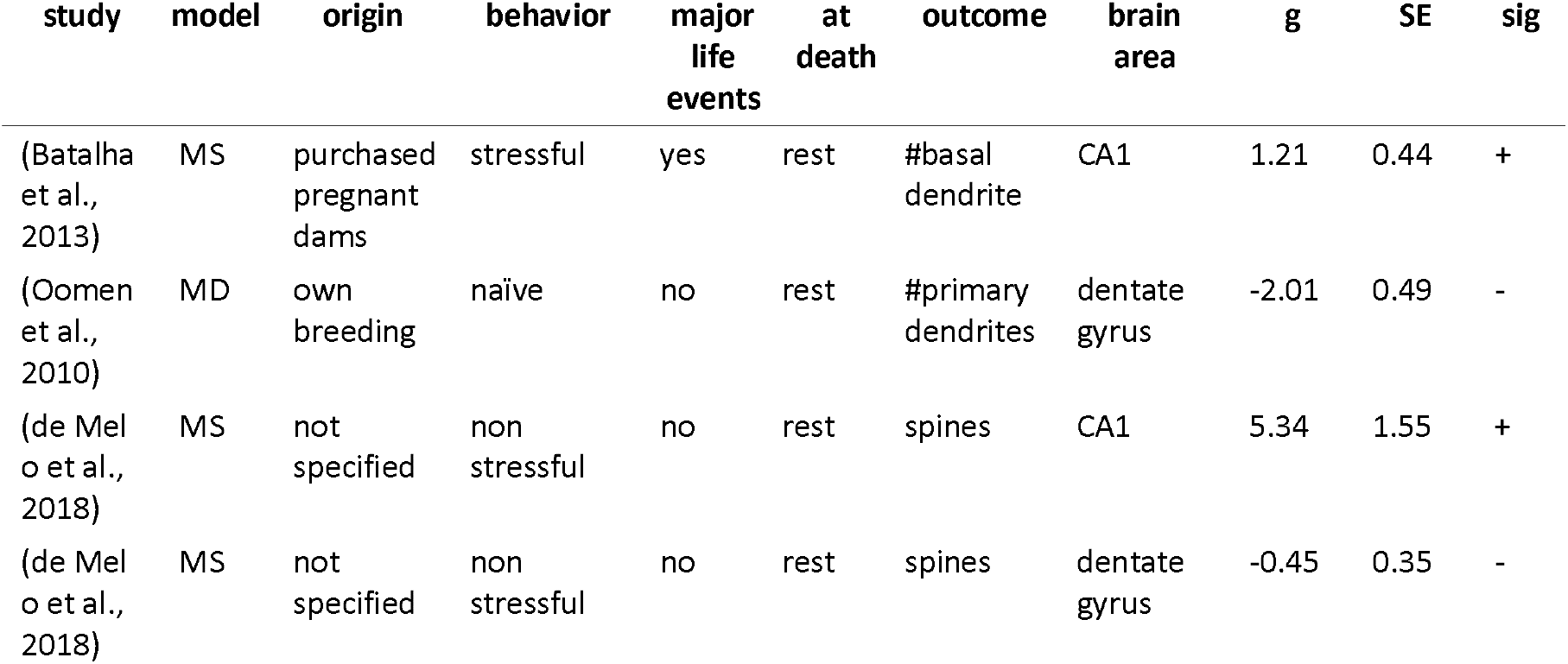
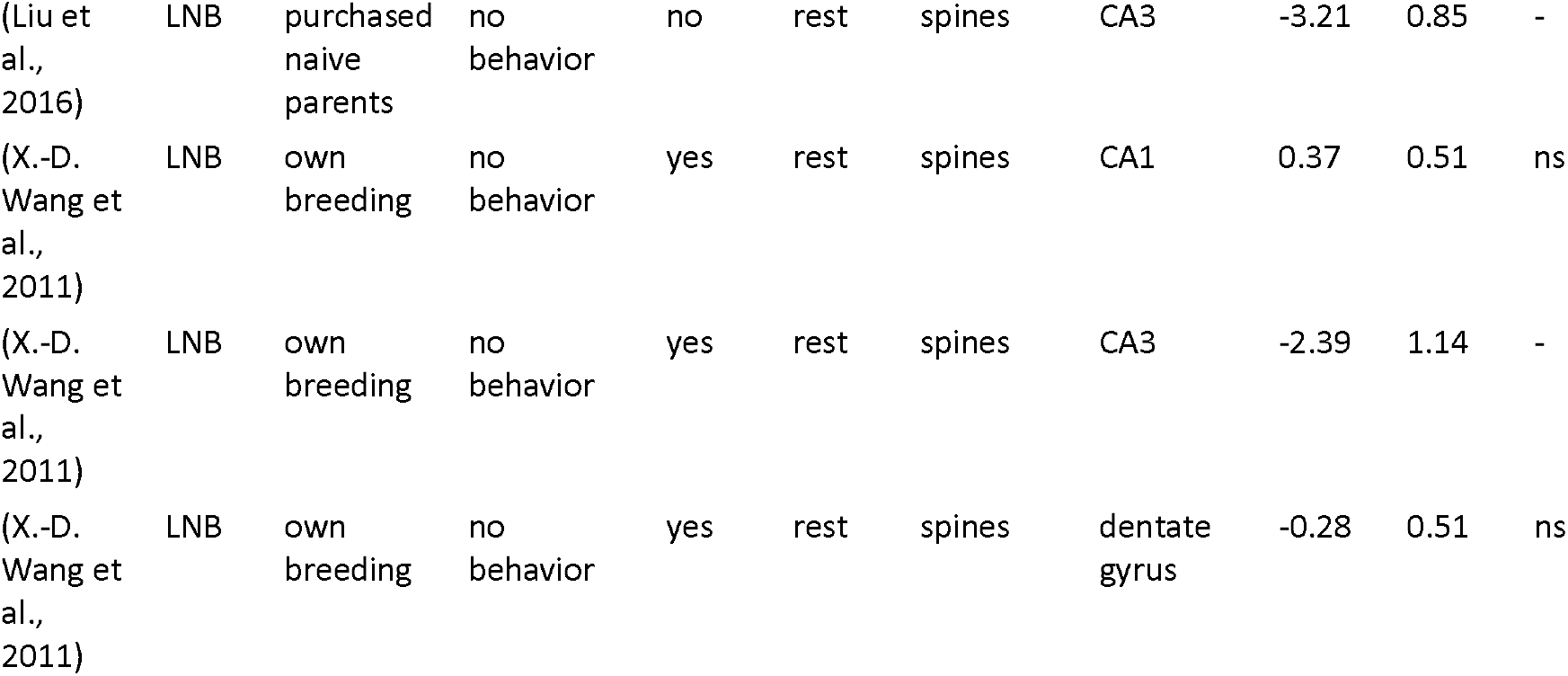
Summary evidence on spine density and number of dendrites. MS = Maternal Separation, MD = Maternal Deprivation, LNB = limited nesting and bedding; in ‘Origin’, we consider purchasing pregnant dams as a stressful experience due to transportation stress of the dams; in ‘Behavior’, naïve = animals left undisturbed (regardless of ELA or control treatment), non stressful = animals that performed non-stressful behavior tests (e.g. open field), stressful = animals that performed stressful behavior tests (e.g. fear conditioning), no behavior = the animals were not naïve, but did not perform behavioral tests (e.g. saline injections); in ‘Major Life Events’, experiments score “yes” if the animals experienced other (besides ELA, prenatal transport stress and stressful behavior tests) traumatic life events e.g. chronic stress during adolescence; in ‘at death’, we defined the status of the animal at death, i.e. at rest or aroused/stressed; g = Hedge’s g; SE = Sampling Error; sig = systematic review significance, where “+” means increase, “-” means decrease and “ns” means “not significant”.

| study | model | origin | behavior | major<br>life<br>events | at<br>death | outcome | brain<br>area | g | SE | sig |
| --- | --- | --- | --- | --- | --- | --- | --- | --- | --- | --- |
| (Batalha et al., 2013) | MS | purchased pregnant dams | stressful | yes | rest | #basal dendrite | CA1 | 1.21 | 0.44 | + |
| (Oomen et al., 2010) | MD | own breeding | naïve | no | rest | #primary dendrites | dentate gyrus | -2.01 | 0.49 | - |
| (de Melo et al., 2018) | MS | not specified | non stressful | no | rest | spines | CA1 | 5.34 | 1.55 | + |
| (de Melo et al., 2018) | MS | not specified | non stressful | no | rest | spines | dentate gyrus | -0.45 | 0.35 | - |
| (Liu et al., 2016) | LNB | purchased naive parents | no behavior | no | rest | spines | CA3 | -3.21 | 0.85 | - |
| (X.-D. Wang et al., 2011) | LNB | own breeding | no behavior | yes | rest | spines | CA1 | 0.37 | 0.51 | ns |
| (X.-D. Wang et al., 2011) | LNB | own breeding | no behavior | yes | rest | spines | CA3 | -2.39 | 1.14 | - |
| (X.-D. Wang et al., 2011) | LNB | own breeding | no behavior | yes | rest | spines | dentate gyrus | -0.28 | 0.51 | ns |

### Quantitative analysis of morphology

A 3-level model was used to investigate whether ELA significantly impacted morphology of the adult hippocampus. In particular, we analyzed i) whether the effects differed across outcomes (volume of the brain area, number of dendritic branches and length of dendrites); and ii) whether other traumatic life experiences interacted with the effects. Of note, the groups compared always differed only in the experience (or not) of ELA. Therefore, effects of other traumatic life events should be considered as “enhancing” (or not) the effects of early life adversity.

Overall, ELA significantly reduced the volume of the hippocampus (*g*(se) = −0.819 (0. 185), t = −4. 424, *p* = 0.001) and decreased the total dendritic length (*g*(se) = −1. 66 (0.303), *t* = −5.473, *p* < 0.001). The decrease in the number of branches per dendritic tree (*g*(se) = −0.699(0.262), *t* = −2.663, *p* = 0.053) was just not significant **(Figure 3a).** The effects were largest when both the ELA and control groups experienced no other traumatic events, (*g*(se) = 1.113(0.268), t = 4.16, *p* = 0.002, **Figure 3c);** qualitatively, this was consistent across all outcomes (Supplementary Figure 1).

**Figure 3.**
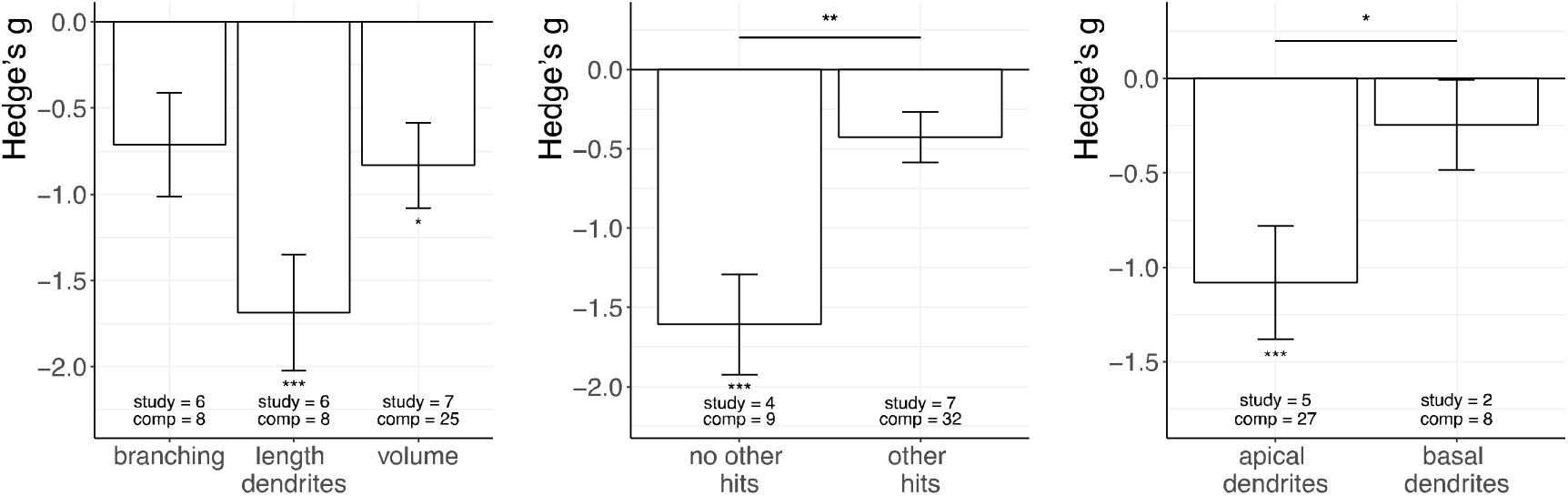
Results on males’ morphology in the hippocampus. A) ELA decreases the volume of the hippocampus and the length of hippocampal dendrites. B) ELA significantly reduces the overall outcome of morphology, both for animals that did not and did experience other major life events. However, the effects are more pronounced when animals (ELA compared to control) did not experience other major life events (“hits”). C) The effects of ELA are more pronounced for apical than for basal dendrites. For each bar the numbers at the bottom refer to the number of studies (study) and comparisons (comp) respectively on which the mean is based. *** = p_adj_ < 0.001; ** = p_adj_ < 0.01; * = p_adj_ < 0.05; study = number of independent publications; comp = number of comparisons (ie difference between ELA and control groups); hits = experience of major life events.

The effects appeared similar in all sub-fields of the hippocampus **(Table 3).** The effects on apical dendrites were more pronounced than on basal dendrites (apical: *g*(se) = −1.08(0.299), *z* = - 3.61, *p* < 0.001; basal: *g*(se) = −0.246 (0.237), *z* = −1.037, *p* = 0.3), as highlighted by a subgroup analysis (*g*(se) = 0.835(0.382), *z* = 2.185, *p* = 0.029, **Figure 3b)**.

**Table 3.** No differences across hippocampal sub-brain areas.

| comparison | g | se | t | p.adj |
| --- | --- | --- | --- | --- |
| dentate vs CA1 + CA3 | 0.158 | 0.312 | 0.508 | 0.875 |
| CA1 vs dentate + CA3 | 0.226 | 0.286 | 0.789 | 0.875 |
| CA3 vs dentate + CA1 | -0.571 | 0.290 | -1.968 | 0.181 |

### Quantitative analysis of neurogenesis

Neurogenesis was determined in the dentate gyrus, more specifically in the subgranular zone; one paper (Bath et al., 2017) was excluded since it reported on the whole hippocampus. Although this may reflect what happens in the dentate gyrus, we excluded the paper to maintain consistency of our sample.

Concerning neurogenesis, ELA significantly decreased the expression of DCX (*g*(se) = −0.825 (0.299), *t* = −2.764, *p* = 0.039), a marker for neuronal differentiation thought to stain immature neurons (Klein et al., 2020) **(Figure 4).** BrdU staining was suppressed after ELA with short (*g*(se) = - 1.335(0.327), *t* = −4.081, *p* = 0.002) but not long (*g*(se) = −1.119(0.435), t = −2.57, *p* = 0.046) delay since injection. These are generally considered as markers of proliferation and survival, respectively (Wojtowicz & Kee, 2006). By contrast, Ki67 expression (a marker of proliferation (Sun & Kaufman, 2018)) was unaffected (*g*(se) = −0.309(0.388), *t* = −0.797, *p* = 0.863). Overall, the results were comparable both for (ELA and control) groups that experienced traumatic life events and those that did not (*g*(se) = 0.182(0.29), *t* = 0.63, *p* = 0.863, **Figure 4b)**.

**Figure 4.**
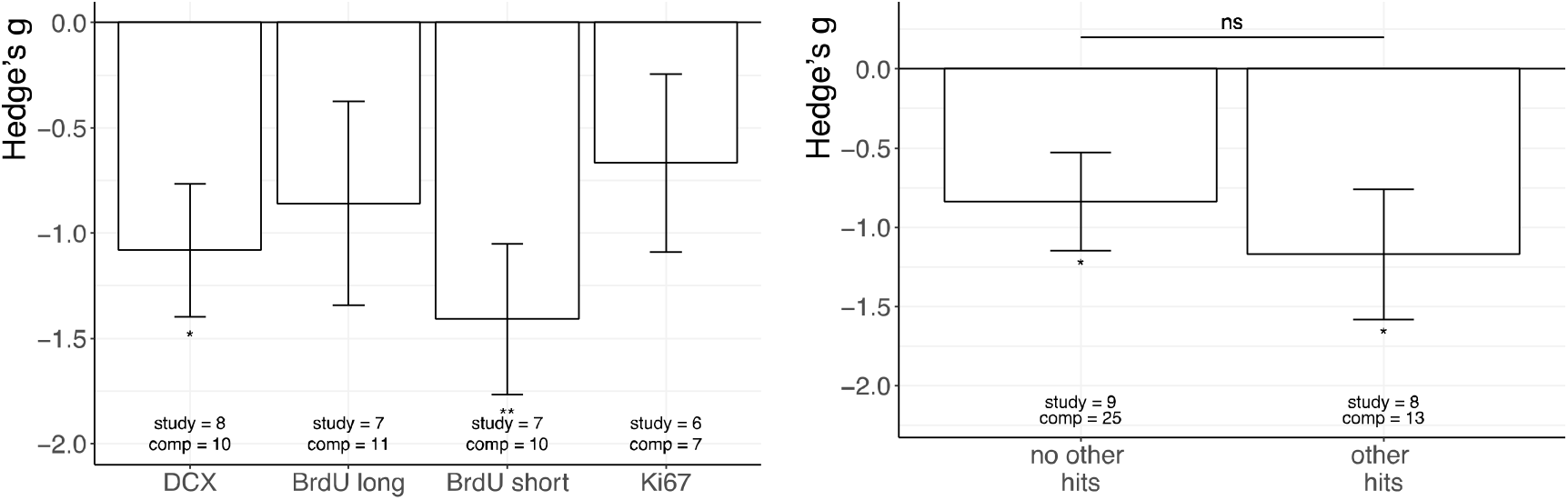
Results of males’ neurogenesis in the hippocampus. A) ELA decreases dcx expression and BrdU expression after a short time since BrdU injection. ELA and control groups are not significantly different in Ki67 expression and in BrdU expression with a long time since injection. B) The effects of ELA are comparable whether or not (ELA and control) animals experienced other major life events (“hits”). *** = p_adj_ < 0.001; ** = p_adj_ < 0.01; * = p_adj_ < 0.05; study = number of independent publications; comp = number of comparisons (ie difference between ELA and control groups); hits = experience of major life events.

### Quantitative analysis of BDNF

Overall, ELA did not alter BDNF expression (*g*(se) = −0.32(0.23), *t* = −1.412, *p* = 0.994), and the pre-specified moderators (type of outcome investigated (mRNA or protein), experience of other traumatic life events, and status of the animal at death (rest / not rest)) did not explain a significant portion of the variance (Q_m_(8) = 8.437, *p* = 0.392). Qualitative exploration of the effect sizes **(Supplementary Figure 2)** suggests that there may be complex 3-way interactions between the factors considered.

### Exploratory analysis with MetaForest

With regard to morphology, in our 3-level model the moderators cumulatively explained a significant portion of the variance (Q_m_(16) = 45.42, *p*<0.001). The remaining heterogeneity was not significant (Q_h_(25) = 21.065, *p* = 0.689, I^2^ = 27.98%), thereby suggesting that no other moderators are necessary to explain the effects of ELA on morphology.

This was different in the case of (all markers of) neurogenesis. Here the moderators did explain a significant portion of the variance (Q_m_(8)=21.762, *p*=0.005); however, there was still unexplained heterogeneity in the model (Q_h_(27)=100.154, *p* < 0.001), suggesting that additional moderators may be relevant to explain the effects. This was next assessed with an exploratory moderator analysis with metaforest. After a thresholded preselection, Metaforest ranks moderators based on how much variance they can explain using random forests. Of the 13 variables investigated, 6 were selected as having a positive variable importance in at least 50% of the 100 bootstrap replications **(Figure 5a).** Specifically, the factor ‘own breeding of the experimental animals’ yielded larger effect sizes compared to animals of different origins (e.g. purchasing naïve parents or pregnant females). Besides this, qualitative exploration of the partial dependency plots **(Supplementary Figure 3)** suggests that the other factors may not be biologically relevant, as supported by the fairly low explained variance (R_cv_^2^(SD) = 0.385(0.33), R_oob_^2^(MSE) = 0.038(1.56)).

**Figure 5.**
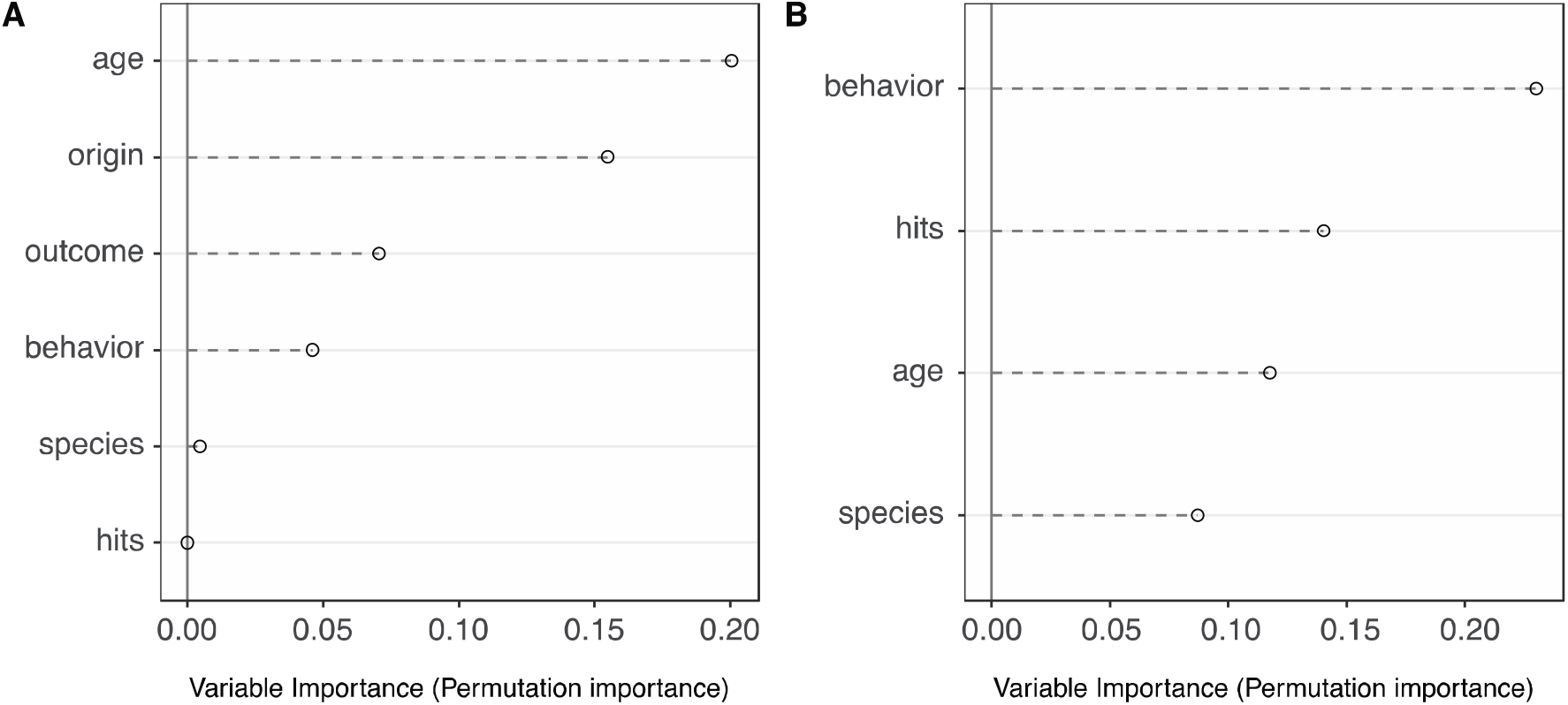
Metaforest variable importance plots after preselection for (A) neurogenesis and (B) BDNF.

The model on BDNF also had moderate remaining unexplained variance (Q_h_(25) = 114.145, *p* < 0.001, I^2^ = 67.33), and from pre-selected confirmatory analysis none of the moderators explained a significant portion of the variance (see section “Quantitative analysis on BDNF”). We therefore chose to use metaforest to explore other potential moderators of the effects. Of the 13 potential moderators selected for metaforest analysis, 3 were selected because they had a positive variable importance in at least 50% of the 100 bootstrap replications **(Figure 5b).** Specifically, partial dependency plots suggest that 1) animals that did not perform behavior tasks and 2) animals that experienced major other life events had larger effect sizes **(Supplementary Figure 4).** However, the explained variance was still low (R_cv_^2^(SD) = 0.366(0.35), R_oob_^2^(MSE) = −0.0256 (1.46)).

### Comparing males and females

As argued, the number of studies reporting on female animals was quite low, prohibiting a full meta-analysis. However, we performed an exploratory analysis in those studies (n_publ_ = 12, n_comp_ = 39) that reported on both sexes. The obtained dataset contained comparisons from all outcomes and brain areas. **Figure 6** visualizes the relationship between male and female data.

**Figure 6.**
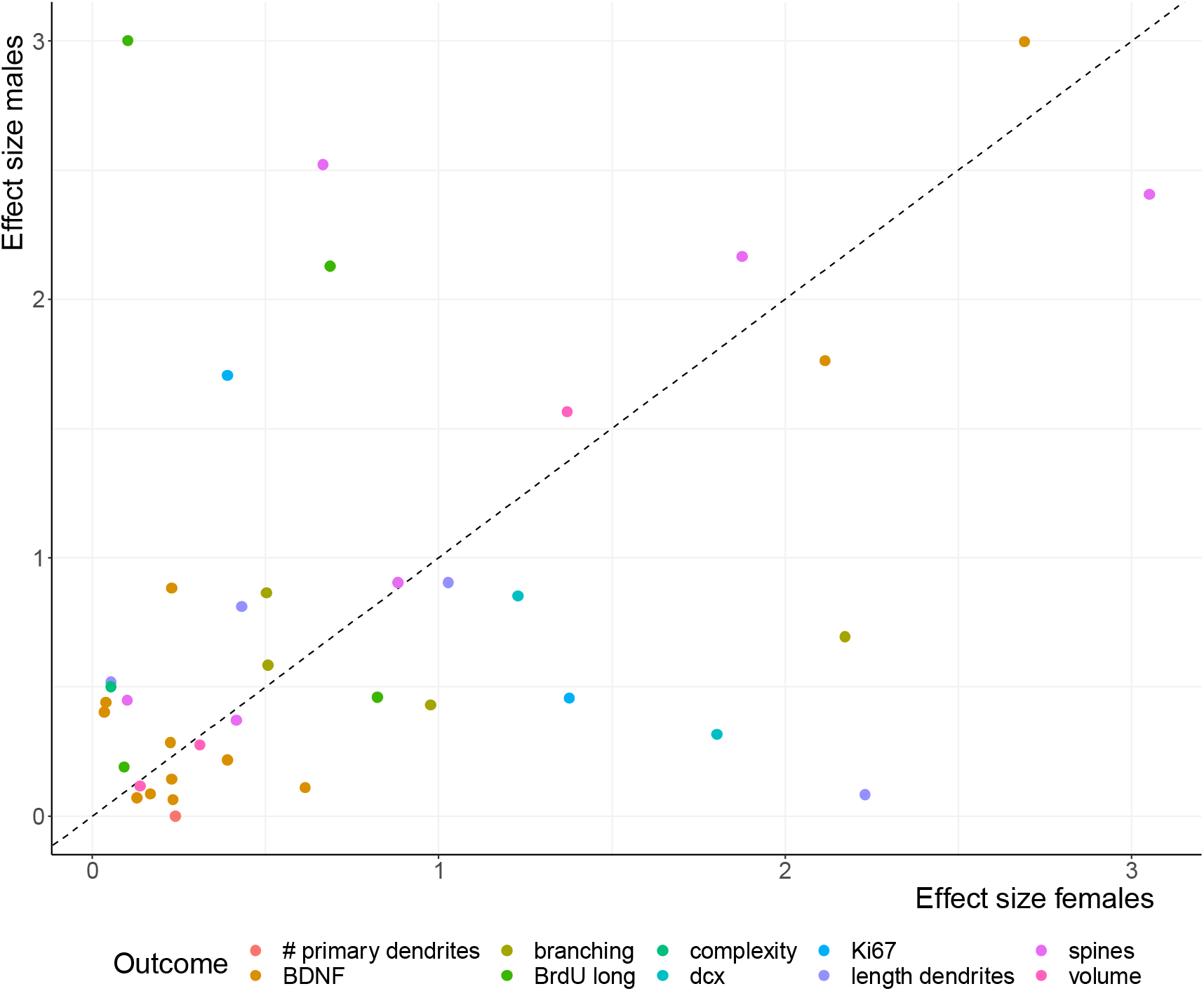
Relationship between male and female effect sizes. Each dot corresponds to the effect size (g) in males and females, obtained in the same publication for the same parameter. The dotted line corresponds to the 45 degrees line where all dots should be if males and females had identical effect sizes. Deviations from the line could be due to error or to real sex differences.

Using a Wald test on this subset of the data no evidence for any sex difference was discerned in effect sizes (*g*_males vs females_ = 0.14(0.14), *z* = 0.996, *p* = 0.32), thereby suggesting that there is no evidence for an overall difference between males and females regarding the effects of ELA on structural plasticity. Given the limited dataset, though, sex differences on specific outcomes and/or brain areas can certainly not be excluded.

### Bias assessment

Risk of bias **(Figure 7)** was assessed using SYRCLE’s risk of bias tool (Hooijmans et al., 2014). Although no publication reported on all items, only two publications did not report being blinded and randomized. However, only four publications provided sufficient information to interpret how randomization was performed. Most importantly, no publication took measures to reduce bias in selective outcome reporting, and this may have been a potential bias in 67.8% of the publications.

**Figure 7.**
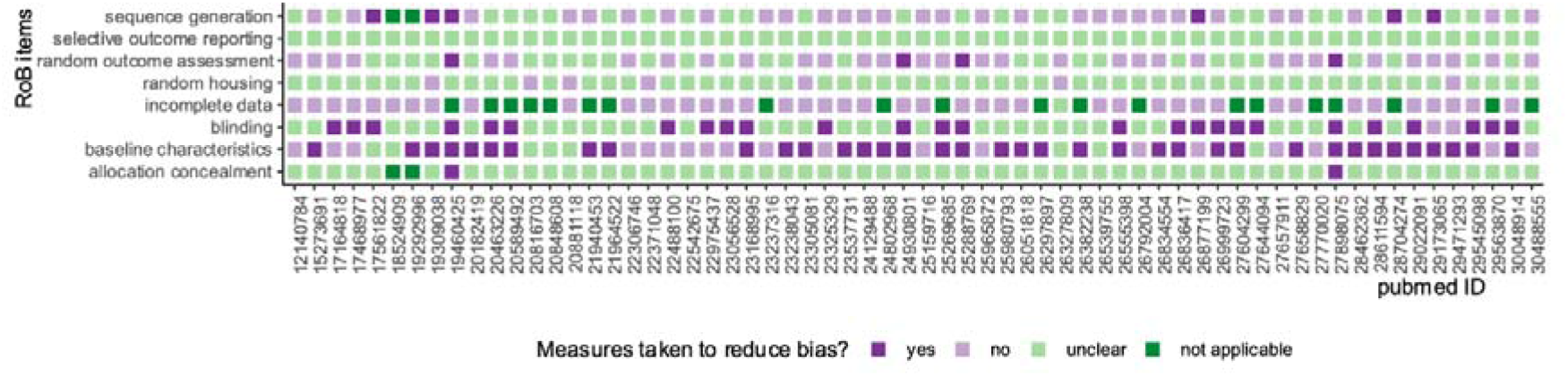
Risk of bias assessment. Each row corresponds to a separate item of the SYRCLE’s risk of bias assessment. Each pubmed ID refers to an independent publication. For the full data, see *https://osf.io/9gru2/*.

We evaluated publication bias by assessing funnel plot asymmetry **(Figure 8)** separately for morphology, neurogenesis and BDNF. Concerning morphology **(Figure 8a)**, there is evidence of asymmetry in the funnel plot. However, only one comparison was present in the highest significance area. Although this may be due to reporting bias, it is unlikely to affect the interpretation of the results. For the neurogenesis and BDNF analyses, the same asymmetry in the funnel plot is observed; yet, here several comparisons are in the high significance area. This suggests that there may be some publication bias, which could lead to an overestimation of effect sizes in the current study. This potential bias could be an important factor when considering the remaining unexplained heterogeneity in our models.

**Figure 8.**
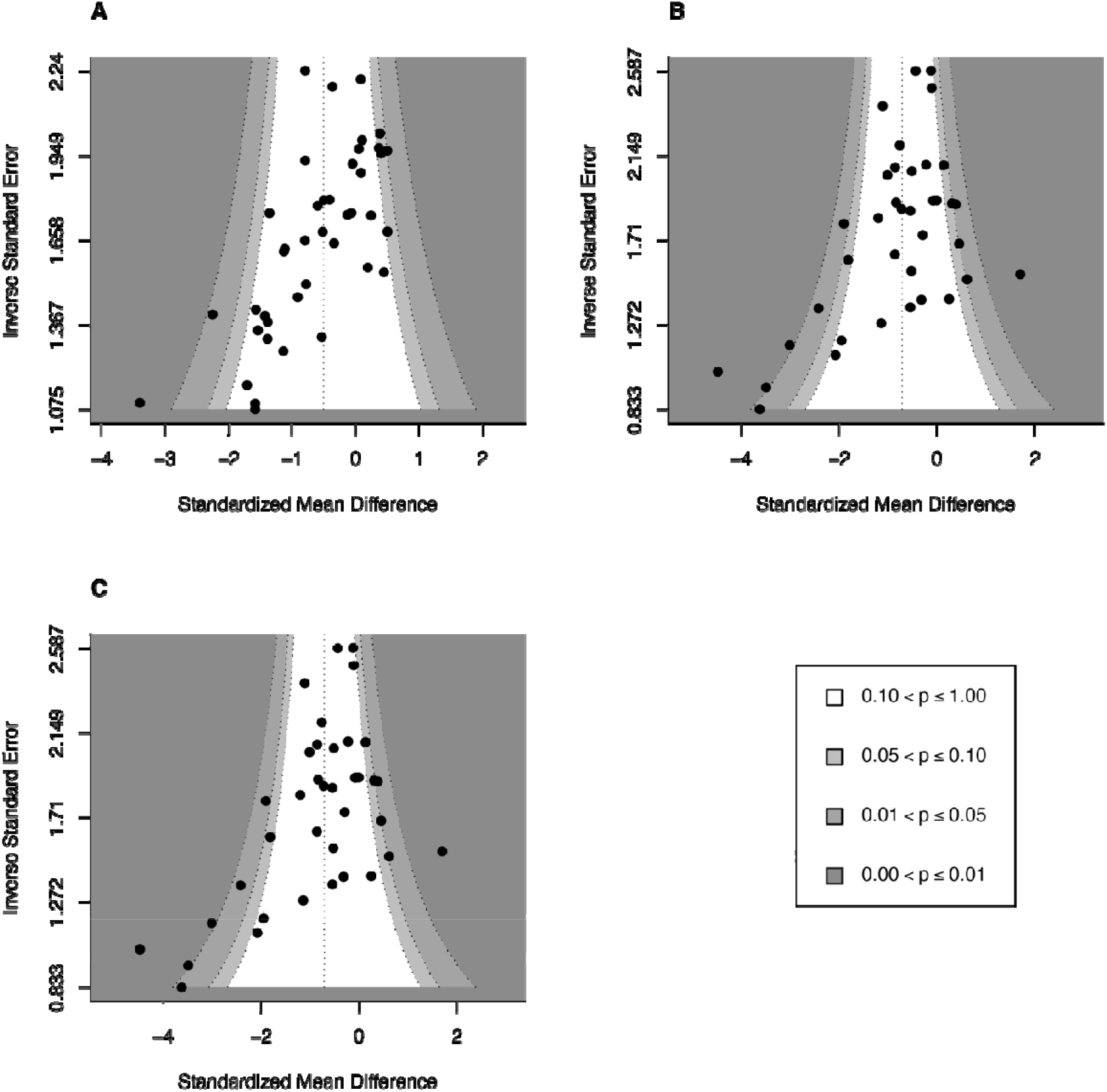
Funnel plot for assessment of publication bias for effects of ELA on (A) morphology, (B) neurogenesis and (C) BDNF analyses. Each dot corresponds to a comparison between control and ELA male groups. *Dark gray = areas of highest significance (p-value). The darker the grey the higher the significance as reported by the legend*.

## Discussion

We set out to review the effects of ELA on neuronal structure and structural plasticity in adult rodents. The survey was restricted to studies in rats and mice, and adversity (here limited to altered maternal care) experienced during the first two weeks of life, focusing on i) the volume of adult brain areas and the morphology of neurons; ii) adult neurogenesis in the dentate gyrus; and iii) expression of BDNF. The descriptive analysis showed that most studies reported on structural changes in the hippocampus of male rats, in most cases using maternal separation as model of ELA. The bias towards the hippocampus is to some extent – but not entirely, i.e. not for morphology – explained by the fact that adult neurogenesis is restricted to a limited number of brain areas, including the dentate gyrus (Ming & Song, 2011). Given the distribution of papers, subsequent quantitative analyses were restricted to observations in the hippocampus of male rodents. Nearly all structural markers were found to be significantly suppressed after ELA compared to control treatment. This was generally not affected by exposure to additional stressors in life or by the state (at rest or aroused / stressed) of the animal immediately before the experiment. Surprisingly, no effects were identified for BDNF, while there was still remaining unexplained heterogeneity. This could possibly be due to data preprocessing (e.g. merging BDNF exons of a subset of papers to maintain consistency across publications), which may have “diluted” the results. Also, we cannot exclude that alterations in BDNF expression took place in the time that elapsed between ELA and the measurements in adulthood. Finally, some caution regarding these conclusions is necessary, since there are indications for publication bias, particularly in the case of neurogenesis and BDNF.

As argued, it is slightly surprising-except in the case of adult neurogenesis-that nearly all studies focused on the hippocampus, with substantially lower numbers of reports on the prefrontal cortex, midbrain dopaminergic areas and the amygdala and (near)absence of studies in the remainder of the brain. It is likely that structural effects of ELA are not restricted to pyramidal neurons in the hippocampus and may well occur e.g. in other pyramidal neurons of the cortex. However, at this stage findings in the hippocampus cannot be simply extrapolated to other regions. This is underlined by, for instance, one group of investigators showing reduced dendritic length in the dentate gyrus of adult male rats earlier exposed to 24 h of maternal deprivation on postnatal day 3 (Oomen et al., 2010), while no such changes were observed in the basolateral amygdala (Krugers et al., 2012). There is a clear need for extension of the current literature to areas other than the hippocampus.

We started out by investigating male and female rodents separately, expecting differences based on earlier reports (Bale & Epperson, 2015; Bangasser & Cuarenta, 2021; Parel & Peña, 2020). The number of reports on female rats or mice, however, was so low that a solid quantitative analysis was not possible. We therefore only carried out an exploratory analysis, using those studies that investigated both sexes. This allowed a comparison in effect sizes in a presumably less heterogeneous sub-group, sharing at least within-study conditions like the experimental procedures, the experimenters carrying out the study and housing conditions of the animals. Although the number of observations was low and varied, there was no evidence that effect sizes were consistently smaller (or larger) in females than in males. Nevertheless, the sparsity of studies in female rodents underlines the message that females are heavily understudied, which may leave potential differences undiscerned (Shansky & Murphy, 2021).

The descriptive analysis also underlined that most studies to date have been carried out in rats rather mice, despite the fact that reliable models for ELA are available in mice too (Peña et al., 2019; Walker et al., 2017). Clearly, there are substantial differences within and between species, e.g. with regard to anxiety-proneness (Clément et al., 2002). To what extent this affects the way in which ELA causes lasting changes in brain structure and structural plasticity remains an unresolved issue until more studies in mice models have been carried out. This also holds true for the type of early life adversity employed in the models, which is currently dominated by maternal separation for several hours during the first 2 postnatal weeks. This model is characterized by a large degree of predictability for the pups (Daskalakis et al., 2011), in contrast to e.g. the limited bedding/ nesting material model (Rice et al., 2008) or a single (24 h) period of maternal deprivation. The latter model has revealed that the exact (postnatal) day of deprivation is crucial for the consequences in adulthood (e.g. (Enthoven et al., 2008)), most likely related to (among other things) the development of the brain and hypothalamus-pituitary-adrenal system at the time of deprivation.

The reduction in volume and dendritic characteristics after ELA were quite robust. The two moderators included in the model, i.e. outcome and cumulative life experiences, explained a significant part of the variation. Of note, we did not consider the possibility that these outcomes would be affected by the state of the animal (at rest versus aroused or stressed) just before the experiment, arguing that changes in volume and particularly dendritic complexity require at least hours to develop. This is different from markers involved in proliferation or the expression of growth factors, which could also be influencing volume. Metaforest analyses suggested that other life experiences too could be important moderators of the effects of ELA on neurogenesis and outcomes. Specifically, origin of the breeding animals and cumulative life experiences were identified for neurogenesis and BDNF, respectively. For BDNF, the presence of other cumulative life experiences in this list may appear as a surprise, since confirmatory moderator analysis was not significant. This is due to the underlying assumptions of the analysis: either due to the selection method (a pre-specified p-value in moderator analysis vs a permutation approach in metaforest), or to non-linear effects that can be established with metaforest but not with the moderator approach. Future research is required to disentangle these two possibilities. Interestingly, origin of the breeding animals and cumulative life experiences (“hits”) were also important moderators for the effects of ELA on behavioral outcomes (Bonapersona et al., 2019). For instance, transporting pregnant dams resulted in stronger effects of ELA on behavioral phenotype than seen with in-house bred dams. In the cases of neurogenesis, also the time elapsed between injecting BrdU and immunohistochemical analysis turned out to be a moderator, likely related to BrdU being an index for proliferation or cell survival, depending on the time elapsed (Wojtowicz & Kee, 2006). Interestingly, while BrdU staining shortly after injection was significantly reduced after ELA, no significant change was observed for the proliferation marker Ki67. One explanation could be that effects of ELA are most apparent in the S-phase of dividing cells (Bannigan, 1985), for which BrdU is a more specific marker than Ki67. Since Ki67 is also present during the G1, G2 and M-phases of cell proliferation, this could have obscured potential effects of ELA in the S-phase.

All in all, we observed a consistent suppressive effect of ELA during the first postnatal weeks on adult structural markers in the hippocampus, specifically on volume, dendritic characteristics and neurogenesis. Possibly, ELA-dependent changes in the activity of growth factors like BDNF could explain such structural changes, although there may be many other pathways through which ELA can lastingly affect structural markers in adulthood. Given the limitation of the vast majority of current reports to the hippocampal area, to one model of early life adversity only (maternal separation) and to male rats, a larger diversity of studies will be necessary to resolve the quest how lasting ELA-dependent structural changes can contribute to changes in behavior.

## Supporting information

Supplementary results and methods

Supplementary Information. Prisma checklist.

## Acknowledgements

We would like to thank Heike Schuler, Dennis van Nuijs, Lieke van Mourik for their help with articles’ selection, and Judith van Luijk for reviewing the study protocol.

## Authors’ contributions

M.J. contributed with the conceptualization, funding acquisition, project administration, supervision, and writing the manuscript; E.K. contributed to the data collection; R.A.S. contributed with the conceptualization, funding acquisition, project administration and supervision; V.B. contributed to the conceptualization, funding acquisition, data collection, data analysis and writing the manuscript.

## Competing interesting

The authors have no competing interests that might be perceived to influence the results and/or discussion reported in this paper.

## Supporting information captions

**S1 File. Supporting information.** Supplementary methods: supplementary note 1 and supplementary table1. Supplementary results: supplementary figures 1 to 4

**S2 File. PRISMA checklist**.

