## Supplementary results and methods for "Structural changes after early life adversity in rodents: a systematic review with meta-analysis"

Marian Joëls^1,2^, Eline Kraaijenvanger^1^, R. Angela Sarabdjitsingh^1^ and Valeria Bonapersona^1, *^

^1^ Department of Translational Neuroscience, UMC Utrecht Brain Center, Utrecht University, The Netherlands

^2^ University of Groningen, University Medical Center Groningen, The Netherlands

* Corresponding Author

Email: v.bonapersona-2 (at) umcutrecht.nl (VB)

This document contains:

Supplementary note 1

Supplementary table 1

Supplementary figures 1-4

Data, scripts and other materials are available at: *https://osf.io/9gru2/*

### Supplementary Notes

#### Supplementary Note 1: search string

**Pubmed:**

*Part 1 - Mice and rats:*

("rodentia"[Mesh] OR rodent*[tiab] OR "mus"[Tiab] OR "mice"[Mesh] OR "mice"[tiab] OR "mouse"[tiab] OR "rats"[Mesh] OR "rats"[tiab] OR "rat"[tiab])

*Part 2 – Postnatal early-life adversity:*

("maternal behavior"[MeSh] OR "maternal care”[tiab] OR "early life stress"[tiab] OR "ELS"[tiab] OR "early life adversity"[tiab] OR "early life adversities"[tiab] OR "ELA"[tiab] OR "early life manipulation"[tiab] OR "early life manipulations"[tiab] OR "early adverse experience"[tiab] OR "early adverse experiences"[tiab] OR "early adversed experience"[tiab] OR "early adversed experiences"[tiab] OR "perinatal stress"[tiab] OR "perinatal adversity"[tiab] OR "perinatal adversities"[tiab] OR "perinatal manipulation"[tiab] OR "perinatal manipulations"[tiab] OR "perinatal adverse experience"[tiab] OR "perinatal adverse experiences"[tiab] OR "perinatal adversed experience"[tiab] OR "perinatal adversed experiences"[tiab] OR "postnatal stress"[tiab] OR "postnatal adversity"[tiab] OR "postnatal adversities"[tiab] OR "postnatal manipulation"[tiab] OR "postnatal manipulations"[tiab] OR "postnatal adverse experience"[tiab] OR "postnatal adverse experiences"[tiab] OR "postnatal adversed experience"[tiab] OR "postnatal adversed experiences"[tiab] OR "neonatal stress"[tiab] OR "neonatal adversity"[tiab] OR "neonatal adversities"[tiab] OR "neonatal manipulation"[tiab] OR "neonatal manipulations"[tiab] OR "neonatal adverse experience"[tiab] OR "neonatal adverse experiences"[tiab] OR "neonatal adversed experience"[tiab] OR "neonatal adversed experiences"[tiab] OR "Maternal Deprivation"[Mesh] OR "maternal deprivation”[tiab]OR "maternal separation”[tiab] OR "limited bedding”[tiab] OR "limited nesting”[tiab] OR "limited material”[tiab] OR "limited bedding/nesting"[tiab] OR "limited bedding-and-nesting"[tiab] OR "limited nesting/bedding"[tiab] OR "limited nesting-and-bedding"[tiab] OR "early life isolation"[tiab] OR "perinatal isolation"[tiab] OR "postnatal isolation"[tiab] OR "neonatal isolation"[tiab] OR "licking and grooming"[tiab] OR "licking-and-grooming"[tiab] OR "licking/grooming"[tiab] OR "early handling"[tiab] OR "early life handling"[tiab] OR "perinatal handling"[tiab] OR "postnatal handling"[tiab] OR "neonatal handling"[tiab])

**Embase search string**

*Part 1 – Mice and rats:*

(rodent*:ab,ti OR mus:ab,ti OR mouse:ab,ti OR mice:ab,ti OR rat:ab,ti OR rats:ab,ti)

*Part 2 – Postnatal early-life adversity:*

('maternal behavior':ab,ti OR 'maternal care':ab,ti OR 'early life stress':ab,ti OR 'els':ab,ti OR 'early life adversity':ab,ti OR 'early life adversities':ab,ti OR 'ela':ab,ti OR 'early life manipulation':ab,ti OR 'early life manipulations':ab,ti OR 'early adverse experience':ab,ti OR 'early adverse experiences':ab,ti OR 'early adversed experience':ab,ti OR 'early adversed experiences':ab,ti OR 'perinatal stress':ab,ti OR 'perinatal adversity':ab,ti OR 'perinatal adversities':ab,ti OR 'perinatal manipulation':ab,ti OR 'perinatal manipulations':ab,ti OR 'perinatal adverse experience':ab,ti OR 'perinatal adverse experiences':ab,ti OR 'perinatal adversed experience':ab,ti OR 'perinatal adversed experiences':ab,ti OR 'postnatal stress':ab,ti OR 'postnatal adversity':ab,ti OR 'postnatal adversities':ab,ti OR 'postnatal manipulation':ab,ti OR 'postnatal manipulations':ab,ti OR 'postnatal adverse experience':ab,ti OR 'postnatal adverse experiences':ab,ti OR 'postnatal adversed experience':ab,ti OR 'postnatal adversed experiences':ab,ti OR 'neonatal stress':ab,ti OR 'neonatal adversity':ab,ti OR 'neonatal adversities':ab,ti OR 'neonatal manipulation':ab,ti OR 'neonatal manipulations':ab,ti OR 'neonatal adverse experience':ab,ti OR 'neonatal adverse experiences':ab,ti OR 'neonatal adversed experience':ab,ti OR 'neonatal adversed experiences':ab,ti OR 'maternal deprivation':ab,ti OR 'maternal separation':ab,ti OR 'limited bedding':ab,ti OR 'limited nesting':ab,ti OR 'limited material':ab,ti OR 'limited bedding/nesting':ab,ti OR 'limited bedding-and-nesting':ab,ti OR 'limited nesting/bedding':ab,ti OR 'limited nesting-and-bedding':ab,ti OR 'early life isolation':ab,ti OR 'perinatal isolation':ab,ti OR 'postnatal isolation':ab,ti OR 'neonatal isolation':ab,ti OR 'licking and grooming':ab,ti OR 'licking-and-grooming':ab,ti OR 'licking/grooming':ab,ti OR 'early handling':ab,ti OR 'early life handling':ab,ti OR 'perinatal handling':ab,ti OR 'postnatal handling':ab,ti OR 'neonatal handling':ab,ti)

### Supplementary Tables

#### Supplementary table 1: model building considerations

Prior the beginning of the study, we identified several factors that may be important moderators of the effects of ELA on structural plasticity, namely: 1) specific outcome parameters, 2) brain area(s), 3) experience of other traumatic events, 4) product measured (mRNA or protein, only for the outcome BDNF), 5) state of the animal at death (only for BDNF and neurogenesis), 6) delay between the start of the experimental manipulation and measuring the outcome (only for the neurogenesis-related parameter brdu). Prior the beginning of the study, we chose that these moderators needed to be addressed in our study, as moderators, filtering variables, subgroup analysis or sensitivity analysis. The choice was based on the distribution of the factors of these variables in the dataset, so that to maximize interpretability (max interaction of 3 moderators) and to minimize the number of tests used. Specific outcome parameters and brain areas were important variables used for filtering. In particular, we performed out analysis only on the hippocampus, because it was the brain area most investigated. We excluded from the quantitative synthesis those outcomes reported by a limited number of publications (for specifics, see table below). With this filtering, we were able to maximize the homogeneity of our dataset for quantitative analysis. The table below reports the analytical considerations for each of the final models.

| **Structural plasticity:** | **Final model** | **Excluded data (filtering)** | **Considerations** |
| --- | --- | --- | --- |
| **Morphology** | Interaction between sub-part of the hippocampus and experience of other traumatic events | Complexity as a composite measure (n_paper_ = 1), spine density (n_paper_ = 3, n_comp_ = 6), number of dendrites (n_paper_ = 3, n_comp_ = 6) |  |
| **Neurogenesis** | Experience of other traumatic events. Sub-parts of hippocampus not applicable (only dentate gyrus included). Subgroup analysis for brdu with short induction and ki67 for interaction of state of the animals at death. | Data not specific to the dentate gyrus (n_paper_ = 1) | Brdu with short (<1 day) vs long (> 1 day) induction time are considered two separate outcomes to decrease the number of moderators used. |
| **BDNF** | Interaction between other traumatic events, state of the animal at death and product measured. These moderators were selected because they explained a significant proportion of the variance in univariate models. |  | State of the animal at death is relevant only for RNA outcomes because for most studies there was a short interval between the induction of the arousal/stress state and decapitation. Sub-parts of the hippocampus were not included in the final model because the majority of observations were measured in the whole hippocampus (58.3% of BDNF comparisons). |

### Supplementary Figures

#### Supplementary Figure 1 - Morphology

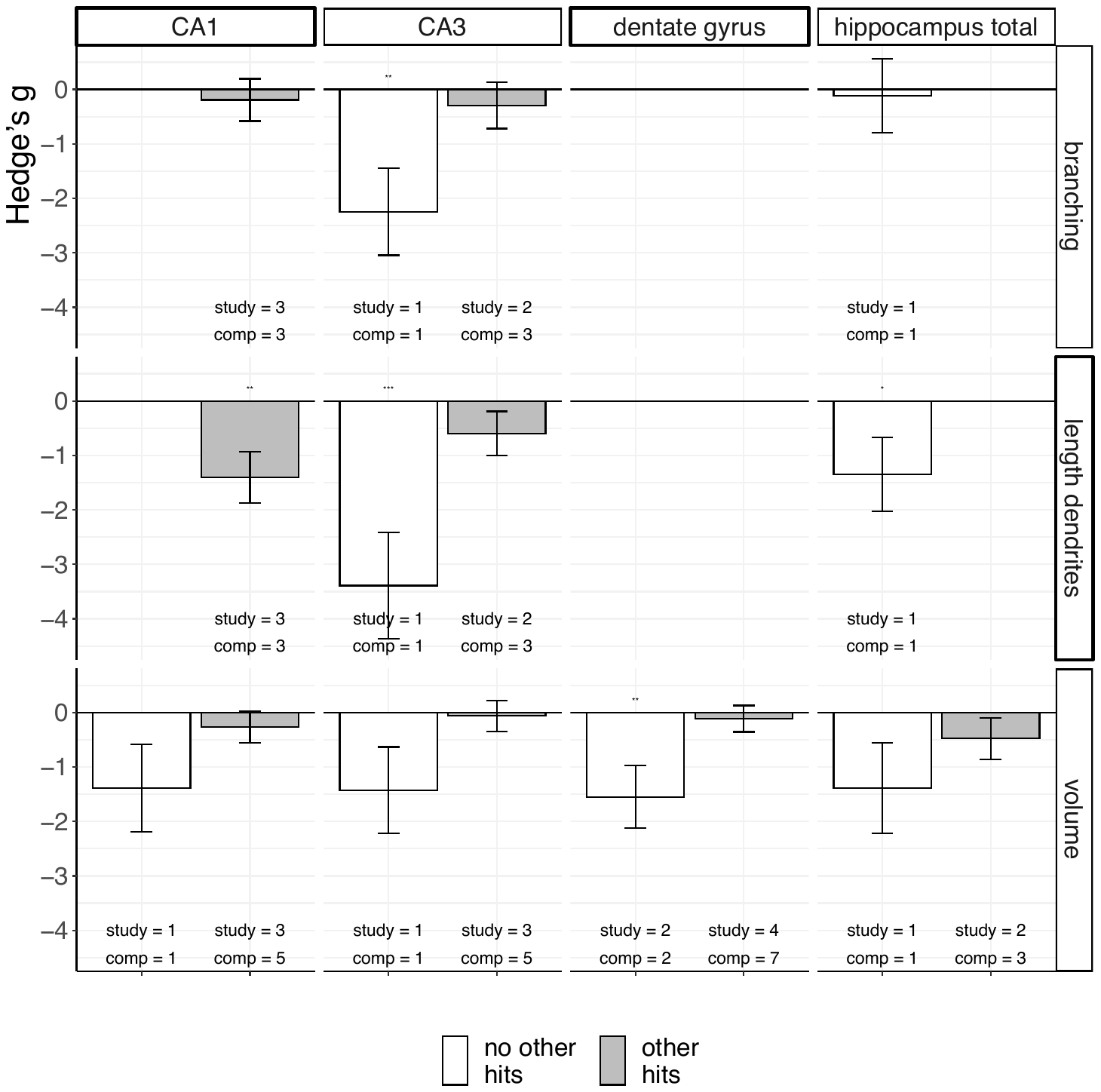

Effect estimates of morphology divided by sub-part of the hippocampus (vertical facets), outcome (horizontal facets) and the experience of additional life traumas (white and grey bars, see legend). *Study = # of publications for the specific outcome; comp = # of comparisons of the specified outcome. * = p < 0.05; ** = p < 0.01; *** = p < 0.001. The p-values reported are not adjusted and should be interpreted as exploratory only.*

#### Supplementary Figure 2 – BDNF analysis

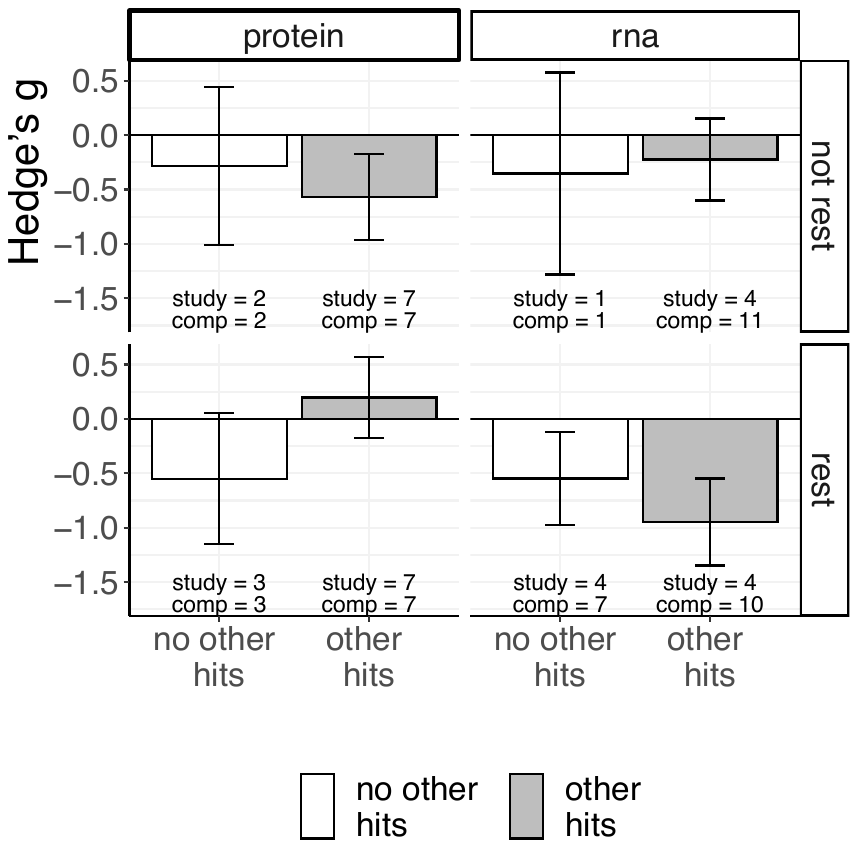

Effect estimates of BDNF divided by RNA/protein (vertical facets), state of the animal at death (horizontal facets) and the experience of additional life traumas (white and grey bars, see legend). *Study = # of publications for the specific outcome; comp = # of comparisons of the specified outcome. * = p < 0.05. The p-values reported are not adjusted and should be interpreted as exploratory only.*

#### Supplementary Figure 3

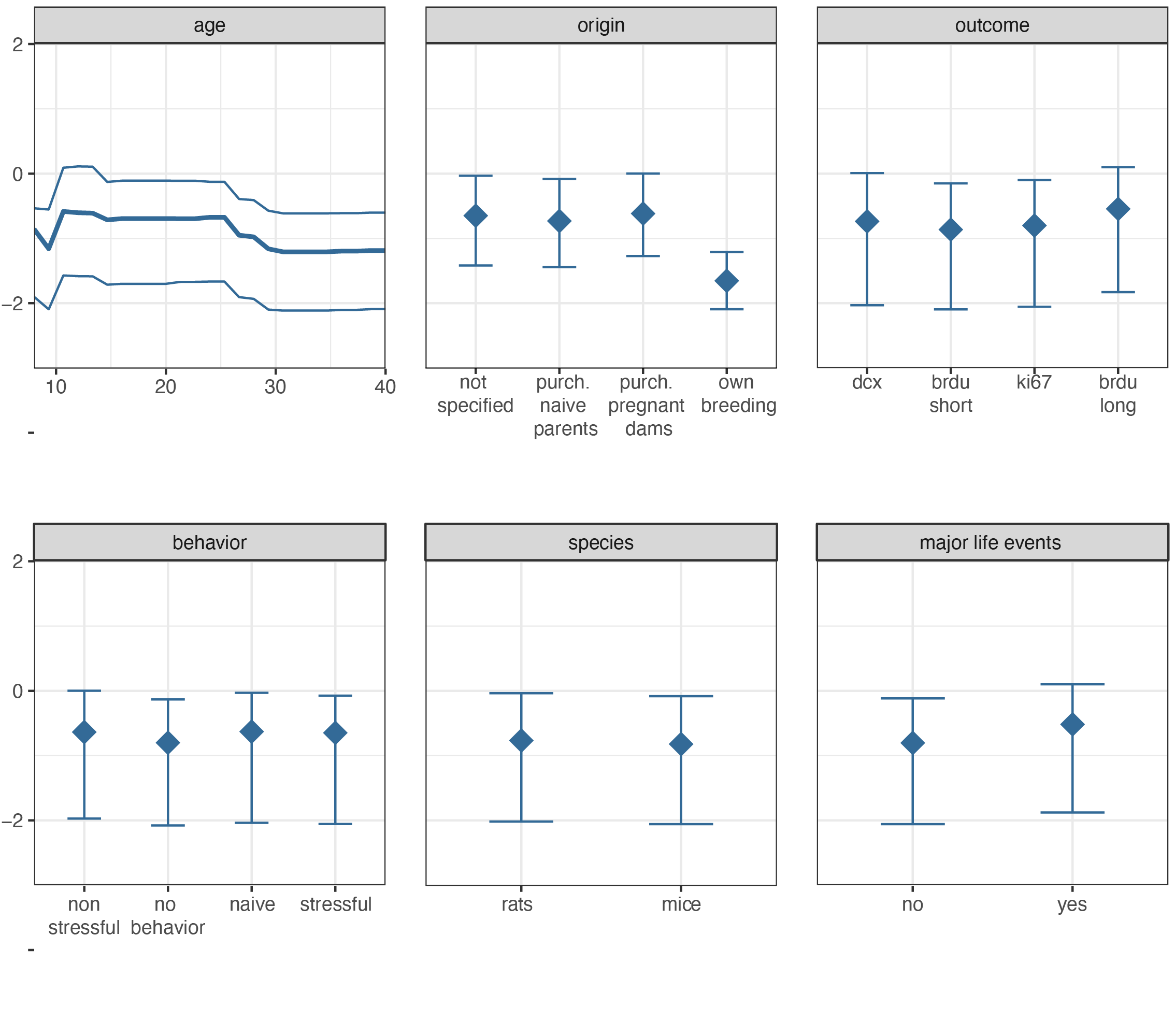

Neurogenesis partial dependency plots of those variables that in at least 50% of the replications had a positive variable importance (MetaForest analysis). Interval corresponds to prediction interval. The changes between sub-groups of each factor appear minor, with exception of origin “own breeding”.

#### Supplementary Figure 4

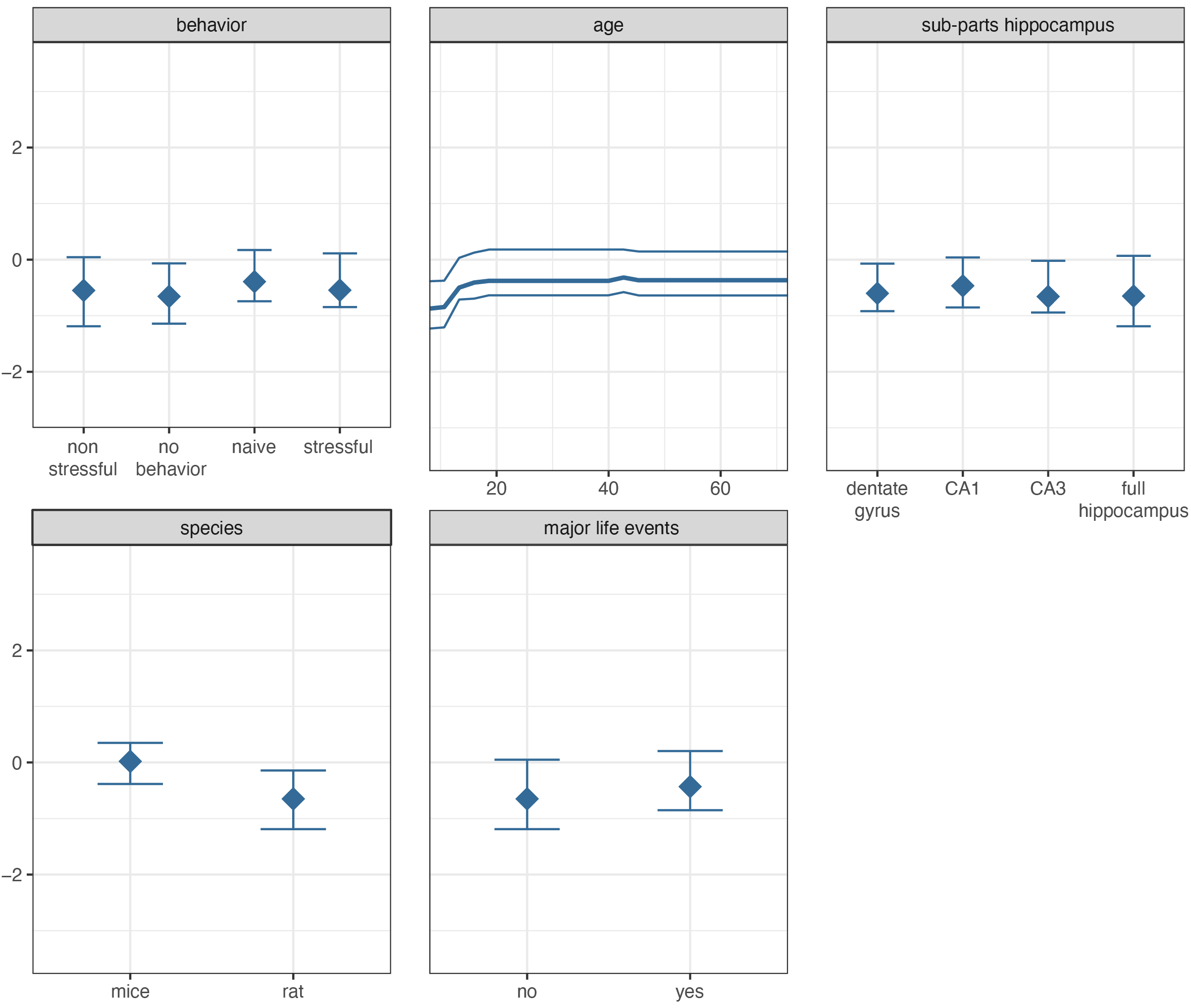

BDNF partial dependency plots of those variables that in at least 50% of the replications had a positive variable importance (MetaForest analysis). Interval corresponds to prediction interval. The changes between sub-groups of each factor appear minor.
