## Supplementary Information. Prisma checklist. for "Structural changes after early life adversity in rodents: a systematic review with meta-analysis"

### PRISMA 2020 Main Checklist

| **Topic** | **No.** | **Item** | **Location where item is reported** |
| --- | --- | --- | --- |
| **TITLE** |  |  |  |
| **Title** | 1 | Identify the report as a systematic review. | Page 1 |
| **ABSTRACT** |  |  |  |
| **Abstract** | 2 | See the PRISMA 2020 for Abstracts checklist |  |
| **INTRODUCTION** |  |  |  |
| **Rationale** | 3 | Describe the rationale for the review in the context of existing knowledge. | Introduction, paragraph 2 |
| **Objectives** | 4 | Provide an explicit statement of the objective(s) or question(s) the review addresses. | Introduction, line 55 to 63 |
| **METHODS** |  |  |  |
| **Eligibility criteria** | 5 | Specify the inclusion and exclusion criteria for the review and how studies were grouped for the syntheses. | Methods, line 82 to 100 |
| **Information sources** | 6 | Specify all databases, registers, websites, organisations, reference lists and other sources searched or consulted to identify studies. Specify the date when each source was last searched or consulted. | Methods, line 79 to 82 |
| **Search strategy** | 7 | Present the full search strategies for all databases, registers and websites, including any filters and limits used. | Supplementary note 1 |
| **Selection process** | 8 | Specify the methods used to decide whether a study met the inclusion criteria of the review, including how many reviewers screened each record and each report retrieved, whether they worked independently, and if applicable, details of automation tools used in the process. | Methods, line 87 to 89 |
| **Data collection process** | 9 | Specify the methods used to collect data from reports, including how many reviewers collected data from each report, whether they worked independently, any processes for obtaining or confirming data from study investigators, and if applicable, details of automation tools used in the process. | Methods, line 102 to 104 |
| **Data items** | 10a | List and define all outcomes for which data were sought. Specify whether all results that were compatible with each outcome domain in each study were sought (e.g. for all measures, time points, analyses), and if not, the methods used to decide which results to collect. | Methods, line 104 to 124 |
|  | 10b | List and define all other variables for which data were sought (e.g. participant and intervention characteristics, funding sources). Describe any assumptions made about any missing or unclear information. | Methods, line 104 to 124 |
| **Study risk of bias assessment** | 11 | Specify the methods used to assess risk of bias in the included studies, including details of the tool(s) used, how many reviewers assessed each study and whether they worked independently, and if applicable, details of automation tools used in the process. | Methods, line 205 to 2016 |
| **Effect measures** | 12 | Specify for each outcome the effect measure(s) (e.g. risk ratio, mean difference) used in the synthesis or presentation of results. | Methods, line 128 to 131 |
| **Synthesis methods** | 13a | Describe the processes used to decide which studies were eligible for each synthesis (e.g. tabulating the study intervention characteristics and comparing against the planned groups for each synthesis (item 5)). | Methods, line 135 to 137 |
|  | 13b | Describe any methods required to prepare the data for presentation or synthesis, such as handling of missing summary statistics, or data conversions. | Methods, line 147-162 |
|  | 13c | Describe any methods used to tabulate or visually display results of individual studies and syntheses. | n.a. |
|  | 13d | Describe any methods used to synthesize results and provide a rationale for the choice(s). If meta-analysis was performed, describe the model(s), method(s) to identify the presence and extent of statistical heterogeneity, and software package(s) used. | Methods, line 132 to 135, line 219 to 222 |
|  | 13e | Describe any methods used to explore possible causes of heterogeneity among study results (e.g. subgroup analysis, meta-regression). | Methods, line 169 to 186 |
|  | 13f | Describe any sensitivity analyses conducted to assess robustness of the synthesized results. | n.a. |
| **Reporting bias assessment** | 14 | Describe any methods used to assess risk of bias due to missing results in a synthesis (arising from reporting biases). | n.a. |
| **Certainty assessment** | 15 | Describe any methods used to assess certainty (or confidence) in the body of evidence for an outcome. | n.a. |
| **RESULTS** |  |  |  |
| **Study selection** | 16a | Describe the results of the search and selection process, from the number of records identified in the search to the number of studies included in the review, ideally using a flow diagram. | Figure 1 and Results line 226 to 231 |
|  | 16b | Cite studies that might appear to meet the inclusion criteria, but which were excluded, and explain why they were excluded. | n.a. |
| **Study characteristics** | 17 | Cite each included study and present its characteristics. | https://osf.io/9gru2/ |
| **Risk of bias in studies** | 18 | Present assessments of risk of bias for each included study. | Figure 7 |
| **Results of individual studies** | 19 | For all outcomes, present, for each study: (a) summary statistics for each group (where appropriate) and (b) an effect estimate and its precision (e.g. confidence/credible interval), ideally using structured tables or plots. | https://osf.io/9gru2/ |
| **Results of syntheses** | 20a | For each synthesis, briefly summarise the characteristics and risk of bias among contributing studies. | n.a. |
|  | 20b | Present results of all statistical syntheses conducted. If meta-analysis was done, present for each the summary estimate and its precision (e.g. confidence/credible interval) and measures of statistical heterogeneity. If comparing groups, describe the direction of the effect. | Results' section |
|  | 20c | Present results of all investigations of possible causes of heterogeneity among study results. | Results, section "Exploratory analysis" |
|  | 20d | Present results of all sensitivity analyses conducted to assess the robustness of the synthesized results. | Results, Line 344 to 368 |
| **Reporting biases** | 21 | Present assessments of risk of bias due to missing results (arising from reporting biases) for each synthesis assessed. | Results, Line 230 to 231, 313 to 316 |
| **Certainty of evidence** | 22 | Present assessments of certainty (or confidence) in the body of evidence for each outcome assessed. | Throughout results' section for each outcome |
| **DISCUSSION** |  |  |  |
| **Discussion** | 23a | Provide a general interpretation of the results in the context of other evidence. | Throughout the discussion |
|  | 23b | Discuss any limitations of the evidence included in the review. | Discussion, Line 426 to 428, 437 to 439, 453 to 454, 459 to 461, 465 to 469 |
|  | 23c | Discuss any limitations of the review processes used. | Throughout the discussion |
|  | 23d | Discuss implications of the results for practice, policy, and future research. | Discussion, Line 501 to 508 |
| **OTHER INFORMATION** |  |  |  |
| **Registration and protocol** | 24a | Provide registration information for the review, including register name and registration number, or state that the review was not registered. | Not registered |
|  | 24b | Indicate where the review protocol can be accessed, or state that a protocol was not prepared. | Protocol can be accessed at https://osf.io/9gru2/ |
|  | 24c | Describe and explain any amendments to information provided at registration or in the protocol. | Methods section |
| **Support** | 25 | Describe sources of financial or non-financial support for the review, and the role of the funders or sponsors in the review. | Financial disclosure section |
| **Competing interests** | 26 | Declare any competing interests of review authors. | Competing interests section |
| **Availability of data, code and other materials** | 27 | Report which of the following are publicly available and where they can be found: template data collection forms; data extracted from included studies; data used for all analyses; analytic code; any other materials used in the review. | Methods, Line 222 |

*From:* Page MJ, McKenzie JE, Bossuyt PM, Boutron I, Hoffmann TC, Mulrow CD, et al. The PRISMA 2020 statement: an updated guideline for reporting systematic reviews. MetaArXiv. 2020, September 14. DOI: 10.31222/osf.io/v7gm2. For more information, visit: <www.prisma-statement.org>
